# Large scale interrogation of retinal cell functions by 1-photon light-sheet microscopy

**DOI:** 10.1101/2022.09.26.508527

**Authors:** Suva Roy, Depeng Wang, Andra M. Rudzite, Benjamin Perry, Miranda L. Scalabrino, Mishek Thapa, Yiyang Gong, Alexander Sher, Greg D. Field

## Abstract

Visual processing in the retina depends on the collective activity of large ensembles of neurons organized in different layers. Current techniques for measuring activity of layer-specific neural ensembles rely on expensive pulsed infrared lasers to drive 2-photon activation of calcium-dependent fluorescent reporters. Here, we present a 1-photon light-sheet imaging system that can measure the activity in hundreds of *ex vivo* retinal neurons over a large field of view while simultaneously presenting visual stimuli. This allowed for a reliable functional classification of different retinal ganglion cell types. We also demonstrate that the system has sufficient resolution to image calcium entry at individual synaptic release sites across the axon terminals of dozens of simultaneously imaged bipolar cells. The simple design, a large field of view, and fast image acquisition, make this a powerful system for high-throughput and high-resolution measurements of retinal processing at a fraction of the cost of alternative approaches.

## Introduction

Imaging fluorescence activity of reporters targeted to genetically defined cell types has greatly expanded the kinds of measurements available in neuroscience. For example, calcium imaging allows measuring neural activity across hundreds to thousands of neurons simultaneously^1,2^. It also allows measuring signals at individual synapses and within sub-cellular compartments such as dendrites^3^ without rupturing the cell membrane. However, fluorescence imaging relies on delivering large amounts of light to excite the fluorescent reporter. In the retina, this presents a problem because the excitation light will also drive phototransduction in rod and cone photoreceptors. To overcome this challenge, previous studies have relied on infrared 2-photon excitation of fluorescent reporters^1,4,5^. This greatly reduces photoreceptor activation and allows imaging changes in fluorescence of downstream neurons, while stimulating the photoreceptors in the visible spectrum^6,7^. A drawback of this approach is that it requires femtosecond pulsed infrared lasers, which are expensive to acquire and maintain.

We have developed a 1-photon light-sheet imaging system for *ex vivo* retinal measurements, that has a simple setup and uses a much less expensive laser. The excitation light is restricted to a relatively large (2.25 mm^2^) and thin plane, thus allowing hundreds of retinal neurons to be imaged simultaneously and calcium signals to be resolved at the level of individual synapses. The system leverages the laminar organization of the retina: photoreceptors are in one cellular lamina, retinal interneurons are in a central lamina (containing horizontal, bipolar and amacrine cells), and the retinal ganglion cell (RGC) layer forms a third cellular lamina (containing amacrine and ganglion cells). Between these layers of cell bodies, there are two synaptic laminae: the outer plexiform layer between photoreceptors and interneurons, and the inner plexiform layer (IPL) between interneurons and RGCs^8^. The light-sheet can be directed to a lamina not containing photoreceptors, thereby reducing photoreceptor activation. While a small fraction of the fluorescence excitation light does reach the photoreceptors, it is not efficiently absorbed by the mouse short-wavelength sensitive (S-) opsin^9^. This opsin is expressed at high levels by all cones in the ventral mouse retina. Thus, calcium-dependent changes in fluorescence can be measured in retinal interneurons or RGCs in ventral mouse retina while stimulating the cone photoreceptors with near-ultraviolet light. The mouse has become a major model of visual processing because of its tractable genetics, making this system well-suited to a wide variety of retinal studies^10^.

We demonstrate and validate the utility of the system in two ways. First, we used mice that express Cre recombinase under the control of parvalbumin promoter (PVCre) to express GCaMP6f in a subset of RGCs. We presented a battery of visual stimuli including a full-field amplitude and frequency modulated stimulus (a.k.a. ‘chirp’ stimulus)^1^, moving bars, checkerboard noise, full-field light steps and local bright/dark spots, while imaging calcium-dependent changes in fluorescence. We were able to functionally classify the GCaMP6f expressing RGCs into eight distinct types, consistent with anatomical studies of PV-expressing RGCs^11,12^. Second, we used mice that express Cre recombinase under the control of PCP2 promoter (PCP2Cre) to express GCaMP6f in a subset of bipolar cells (BCs). We were able to resolve hundreds of individual synaptic release sites in BC axons terminals, measure calcium responses to visual stimuli, and functionally classify bipolar cells into an ON (type-6) and an OFF (type-2) type, consistent with previous studies^13^. These results demonstrate that 1-photon light-sheet imaging system is a relatively affordable and viable platform for monitoring and functionally characterizing neural activity across large populations of retinal neurons.

## Results

### 1-photon light-sheet microscope for stimulus delivery and retinal imaging

The microscope system features three main units: (1) a light-sheet fluorescence excitation system, (2) a fluorescence emission detection system, and (3) a visual stimulus delivery system (Fig. 1a). The collimated beam from the laser is transformed by a set of relay lenses and a cylindrical lens to generate a Gaussian light-sheet that is focused by the illumination objective at the position of the sample. The lateral and longitudinal extent of the light-sheet is controlled by a set of apertures. The thickness of the light-sheet is controlled by a horizontal slit and the numerical aperture of the illumination objective. The center of the excitation plane is aligned with the detection axis by adjusting the position of the illumination objective (Fig. 1a).

**Figure 1:**
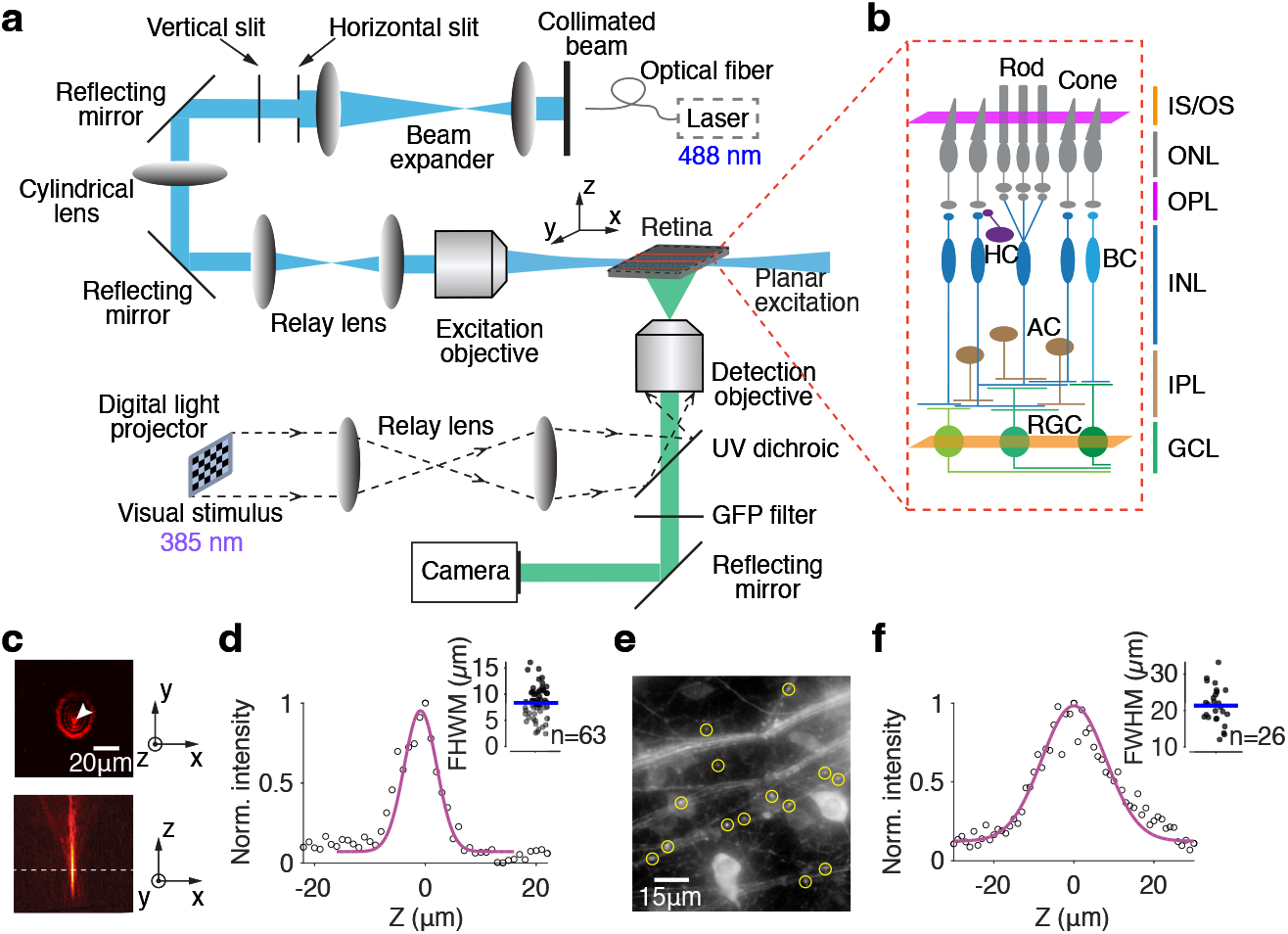
Design and characterization of a 1-photon light-sheet microscope for *ex vivo* retinal imaging. **(a)** Optical arrangement producing a Gaussian beam light-sheet from a 488 nm continuous wave laser source. The light-sheet is coplanar with the retinal imaging plane. The visual stimulus is projected onto the photoreceptors through a detection objective. A high-speed CMOS camera captured the fluorescence images. **(b)** Schematized retinal circuit with stimulus plane (magenta) and imaging plane (orange). IS/OS: inner segment/outer segment of photoreceptors, ONL: outer nuclear layer, OPL: outer plexiform layer, INL: inner nuclear layer, IPL: inner plexiform layer, GCL: ganglion cell layer, HC: horizontal cell, BC: bipolar cell, AC: amacrine cell, RGC: retinal ganglion cell. **(c)** Light-sheet imaging of fluorescent microspheres (500 nm diameter) using a 20x objective. Top: lateral image, bottom: axial image. **(d)** Axial intensity profile of an imaged microsphere. Magenta curve: Gaussian fit. Inset: full-width half max (FWHM) of Gaussian fits for n=63 microspheres. Depth of field = 8.5 ± 2.8 *μ*m. **(e)** GCaMP6f expression in varicosities (yellow circles) along RGC axons. **(f)** Axial intensity profile of a representative varicosite (magenta curve) and depth of field, 22.2 ± 8.1 *μ*m, estimated from n=26 varicosities.

For imaging, the dissected retina was placed inside a chamber with RGC side facing down. The excitation light-sheet was directed parallel to the plane of the retina containing the GCaMP expressing cells. Images were acquired at 10-50 Hz, with a field of view of 700 - 1500 *μ*m along each dimension. Spatial binning was used to improve the signal to noise ratio of the acquired images. An operating laser diode power of 0.1-15 mW, corresponding to 0.01-1.5 mW power at the sample, was found to be adequate for detecting spontaneous and stimulus evoked activity in RGCs.

The visual stimulus for targeting photoreceptors was displayed by a digital light projector (DLP) with a 385 nm LED. The stimulus was delivered to the retina through the objective used for imaging fluorescent emission. To focus the stimulus on the photoreceptor plane without changing the plane of imaging, the position of the DLP relative to the microscope tube lens was adjusted (Fig. 1a, see also Methods). This allowed moving the visual stimulus focal plane by ∼250 *μ*m, sufficient to span the entire thickness of the mouse retina^14^ (Fig. 1b).

### Axial spread of light-sheet from scattering

A key aspect that controls the quality of images of biological tissues is scattering. The more the excitation light is scattered, the higher the background illumination and the worse the image quality. In 1-photon light-sheet imaging, scattering of excitation light can increase the effective thickness of the light-sheet and thus reduce the resolution and image contrast^15^.

To determine the impact of scattering on the thickness of the light-sheet, we first measured the true thickness. This was achieved by imaging 500 nm fluorescent beads embedded in agarose^16^ (Fig. 1c). Agarose has a scattering coefficient of ∼1 cm^-1 17^, and therefore minimally scatters light. The depth of field estimated by fitting the measured axial intensity profile of individual beads with a Gaussian function was 8.5±2.8 *μ*m (Fig. 1d). Next, we modeled the Airy disk profile as a Gaussian with standard deviation equal to the theoretical depth of field, and then deconvolved it from the measured intensity profile to obtain the true thickness of the light-sheet. With a theoretical depth of field 3.48 *μ*m (Eq. 5 in Methods), the true thickness of the light-sheet was 7.75 *μ*m.

The retina has a scattering coefficient of 56.8 cm^-1 18^, meaning that it scatters 1-photon excitation light more than agarose. To determine the true thickness of the 488 nm light-sheet in the retina, we imaged individual axon varicosities of RGCs (Fig. 1e). The varicosities are ∼1-4 *μ*m^19^ and exhibit robust calcium fluorescence from action potential induced calcium influx^20^, thus acting as proxies for fluorescent beads. With a measured depth of field ∼22.2±8.1 *μ*m, averaged over n=26 varicosities (Fig. 1f), and using the above deconvolution procedure, the true thickness of the light-sheet was obtained as 21.72 *μ*m. This indicates that the light-sheet, when directed at the ganglion cell layer, and/or the inner plexiform layer, remains well-confined to these laminae^14^ and does not strongly intersect with the photoreceptor layer.

### Targeting S-cones for visual stimulation

A challenge with fluorescence imaging in the retina is that photoreceptors are exquisitely light sensitive, and therefore can be activated by even a small amount of scattered light. The excitation will depend on the overlap between the opsin action spectrum and the excitation wavelength. To overcome this challenge, we targeted our experiments to the ventral mouse retina because the cones in this region predominantly express a short-wavelength sensitive S-opsin that is maximally sensitive to 360 nm (Fig 2b, c)^9,21^. Light sensitivity of this S-opsin at the fluorescent excitation wavelength of 488 nm is ∼4 orders of magnitude lower than its peak sensitivity. This further reduced the effect of light-sheet scatter on cone photoreceptor excitation. Calibration experiments indicated that the rate of S-opsin activations from the scattered light sheet was ∼10^2^ photoisomerizations(P*)/cone/s (Supplementary Fig. 1). By comparison, the visual stimulus focused onto the cones was delivered at an intensity equivalent to ∼10^5^ P*/cone/s, at 385 nm, near the peak S-opsin sensitivity^9,21^. Thus, the scattered light was ∼0.1% of the mean visual stimulus, below the contrast detection sensitivity of cones^22^.

**Figure 2:**
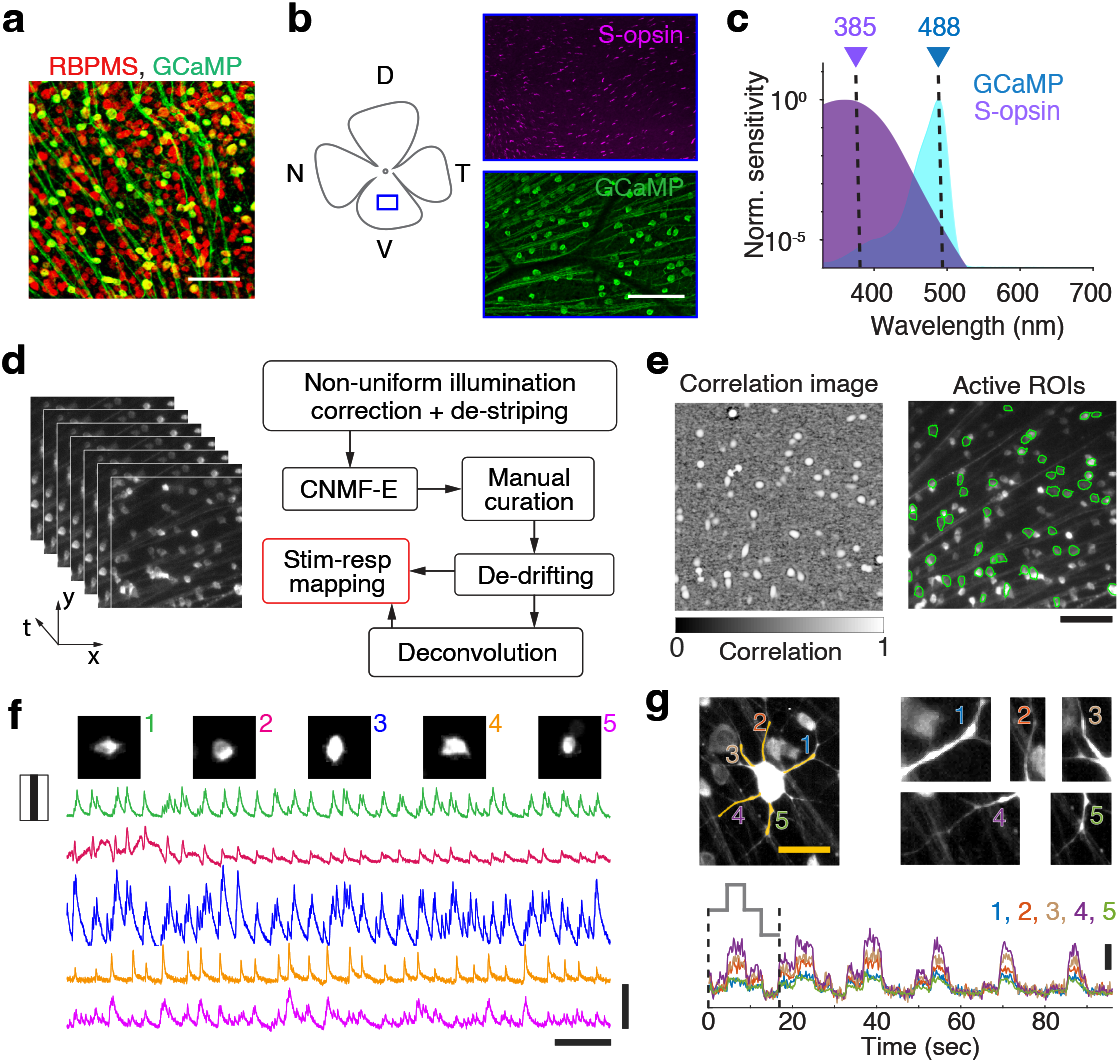
Imaging calcium responses of RGCs. **(a)** RGCs in retina from Ai148;PVCre mouse immunolabeled with pan-RGC marker RBPMS (red), and with GFP (green: GCaMP6f). Scale bar: 50 *μ*m. **(b)** Left: Ventral retina used for fluorescence imaging. Right: Immunolabeling for S-opsin in cone photoreceptors (magenta) and GFP (green: GCaMP6f). Scale bar: 100 *μ*m. **(c)** Visual stimuli delivered at 385 nm (purple arrow) near the peak spectral sensitivity of S-opsin (purple shaded area). Laser excitation at 488 nm (blue arrow) matched to the peak excitation of GCaMP (blue shaded area). **(d)** Image analysis pipeline for extracting spontaneous or visual stimulus evoked calcium responses and inferred spikes of RGCs. **(e)** Left: Spatiotemporal response correlation image. Gray bar: Correlation coefficient. Right: Outlines (green contours) of active RGCs. Scale bar: 100 *μ*m. **(f)** Fluorescent images of representative RGC somata (top) and their temporal responses (bottom) to moving bars. Vertical scale bar: 20 (units of SNR, 6dB cutoff). Horizontal scale bar: 200 s. **(g)** Calcium activity in dendrites of a representative RGC to repeated full field light steps (gray trace, bottom). Horizontal scale bar: 20 *μ*m. Dashed vertical lines indicate duration of a single trial; total n=6 trials. Temporal traces for 5 dendrites (top right) are shown in color. Vertical scale bar: 2 (units of SNR, 6dB cutoff).

### Calcium fluorescence signals in active RGCs

To image calcium activity in RGCs, we used ventral retinas from Ai148;PVCre mice (Fig. 2b) ^23,24^. Robust GCaMP6f fluorescence was observed in the somata, axons and dendrites of RGCs (Fig. 2d-g). To extract and analyze the changes in calcium-dependent fluorescence resulting from visual stimulation of the photoreceptors, fluorescence images were first denoised, corrected for non-uniform illumination, and then processed using a semi-automated algorithm^25^ (see Methods, Fig. 2d). Specifically, a threshold signal to noise ratio^25^ of 8 and a pixel intensity correlation of 0.8 were used to detect and process the signals from fluorescent somata. In a typical experiment, this yielded several hundred active RGCs in the field of view (Fig. 3a). We were also able to measure spontaneous and stimulus driven calcium responses from primary and secondary dendrites of individual RGCs (Fig. 2g). These data confirmed the system’s capability for imaging large-scale neural activity in the retina.

**Figure 3:**
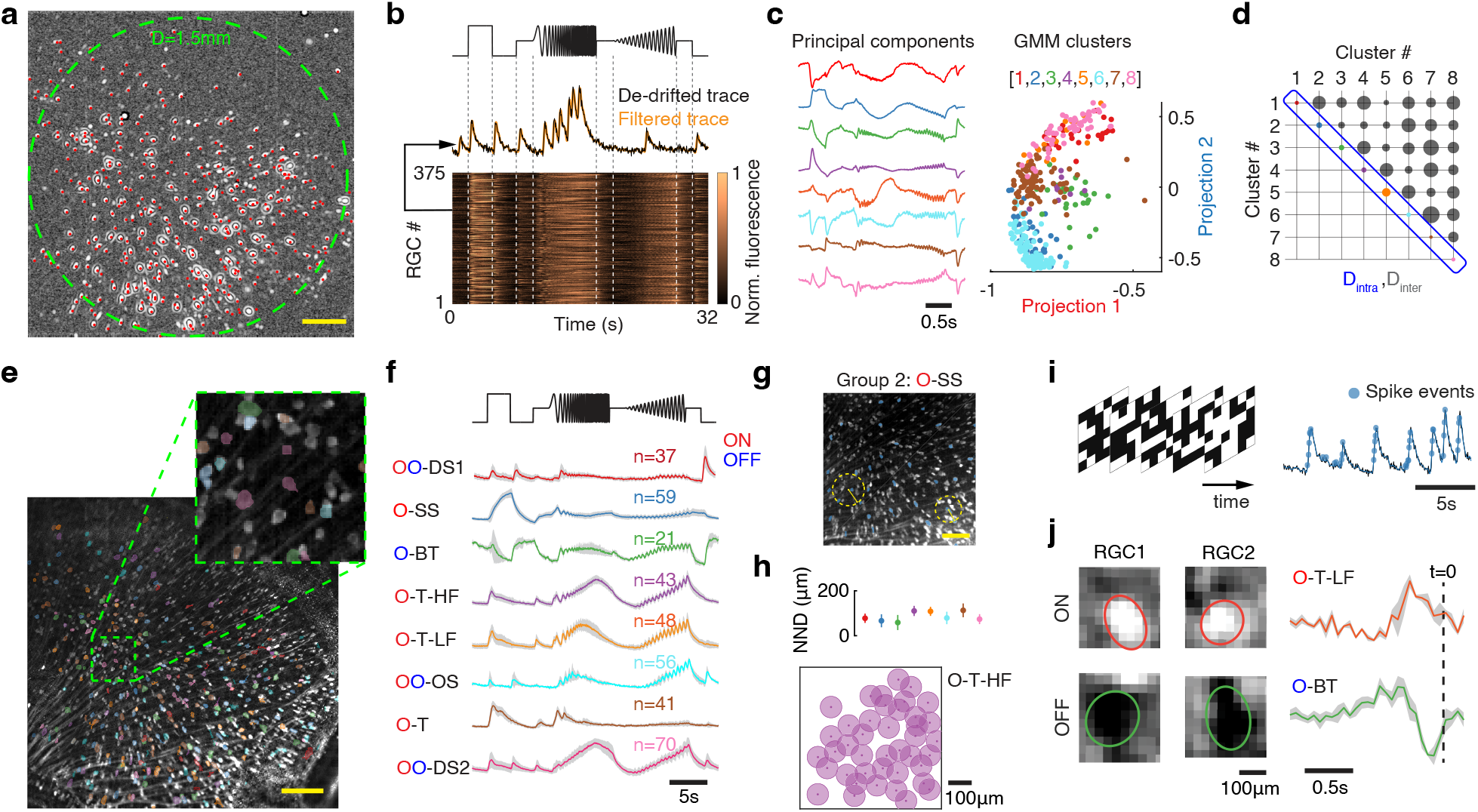
Classification of RGCs from population calcium responses. **(a)** Response correlation image over imaging field of view. Red filled circles: Active RGCs, n=375. Green dashed circle: 1.5 mm diameter. Scale bar: 200 *μ*m. **(b)** Top: Chirp stimulus trace (black). Middle: Baseline corrected (black) and filtered (orange) temporal response of a representative RGC. Bottom: Heatmap of temporal fluorescent traces of n=375 RGCs. **(c)** Left: 8 leading principal components of the normalized temporal responses in (b). Right: Gaussian mixture model (GMM) fit to projection values in 8-D hyperspace shown in a 2-D plane. Clustered projection values (n=8 clusters) are shown by colored circles. **(d)** Mean pairwise Euclidean distance between projection values within each cluster (D_intra_) and between clusters (D_inter_) are illustrated by circles along the diagonal and the off-diagonal respectively. The radius of each circle represents the Euclidean distance. Color scheme for diagonal elements is same as in (c). **(e)** Median intensity projection image with overlaid patches showing soma locations of active RGCs. Patch color corresponds to the clusters in (c). Scale bar: 200 *μ*m. Inset: Magnified view of an image patch. **(f)** Mean temporal responses of RGCs from different clusters to the chirp stimulus (top trace, black). ON: O (red), OFF: O (blue); DS: direction selective; SS: slow sustained; BT: brisk transient; T-HF: transient high frequency; T-LF: transient low frequency; OS: orientation selective; T: transient. Shaded error bar: SD. **(g)** Image showing RGC locations (blue patches) from representative group 2. Dashed circle and box (yellow) depict nearest neighbor RGCs. Scale bar: 100 *μ*m. **(h)** Top: Median (filled circle) and MAD (error bar) of nearest neighbor distances (NND) for each group in (f). Bottom: Representative mosaic of O-T-HF RGCs, based on median NND (radius of circle). **(i)** Checkerboard pattern stimulus (checker size ∼ 40 *μ*m), corresponding response trace of a representative RGC (black) and inferred spikes (light blue circles). **(j)** Spatial RFs of representative RGCs (left) and mean temporal RFs across all cells (right) belonging to ON transient low frequency and OFF brisk transient types. Time of spike: t=0. Shaded error bar: SD.

### Response classification of RGCs using full-field chirp and checkerboard noise stimuli

To validate measurement quality in RGC somata, we tested the reliability of functionally classifying RGCs based on the changes in fluorescence produced by visual stimuli. Previous studies show that parvalbumin is expressed in 8 morphologically distinct RGC types^11,12^, suggesting an equivalent number of functionally distinct RGC types. To classify the RGCs, we presented a ‘chirp’ visual stimulus^1^ (Fig. 3b). This stimulus consists of a full-field light step, followed by full-field frequency and amplitude modulated sinusoidal illumination. Calcium-dependent fluorescence changes in 375 RGCs (n=1 retina) were acquired over a field of view ∼2.25 mm^2^ (Fig. 3a, b). Singular value decomposition (SVD) was applied to the temporal fluorescence traces and leading principal components were used to generate a projection hyperspace (Fig. 3c, left; also see Methods). Fitting a Gaussian mixture model to the projection space yielded 8 clusters (Fig. 3c, right). The discriminability of the clusters was assessed by comparing the inter-cluster Euclidean distance with intra-cluster Euclidean distance (Fig. 3d). Accuracy of clustering was determined by cross-validating clusters from alternative clustering methods such as Hierarchical agglomerative clustering (HAC)^26^ and Spectral clustering (SC)^27^. Each clustering algorithm produced the same results (Supplementary Fig. 2).

Based on the temporal response properties to different phases of the chirp stimulus, each cluster of RGCs was further assigned to distinct functional types: ON, OFF, ON-OFF, transient and sustained (Fig. 3e, f). The RGCs in each cluster also exhibited a mosaic-like regularity in their spatial arrangement^28^, with nearest neighbor spacing ranging from 70 to 120 *μ*m for different types (Fig. 3g, h). This suggests that each cluster represents a functionally distinct RGC type^29-31^.

To examine the contrast polarity and temporal integration properties of RGCs, we characterized the spatiotemporal receptive fields (RFs) of RGCs from their changes in fluorescence during the presentation of checkerboard noise. Spike trains were estimated from calcium signals, and calcium transients associated with at least 1 spike were used to estimate the RF (see Methods, Fig. 3i). RGCs classified as ON brisk transient and OFF brisk transient types from responses to the chirp stimulus exhibited ON- and OFF-center responses, respectively (Fig. 3j). The temporal responses exhibited positive and negative contrast preferences immediately preceding putative spikes (estimated from the calcium signals), with a biphasic profile consistent with previous findings for these RGC types^28,32^.

### Direction and orientation selective responses of RGCs

To identify direction selective (DS-) RGCs, we imaged calcium responses of RGCs from the same retina as in Figure 3, to bright bars (100% contrast) moving along 12 different directions. We calculated a direction selective index (DSI) using the area under the temporal fluorescence trace for different movement directions (see Methods). RGCs with DSI greater than 0.3 were classified as DS-RGCs^33,34^ (Fig. 4a, b, d). Prolonged exposure to excitation light can bleach GCaMP, thereby reducing its sensitivity over time^6^. This can bias DSI estimates obtained from trial averaged responses. Therefore, only trial blocks in which the DSI did not change by more than 20% of that estimated from the first trial, were included in the analysis. A total of 29 RGCs were identified as direction selective (Fig. 4d), out of which 21 had clear ON and OFF responses (Fig. 4b inset). The ON-OFF DS-RGCs were further classified into 4 subtypes^35^ based on orthogonality of preferred directions (Fig. 4e). A similar fraction of DS-RGCs (∼10%) were estimated from retinas of other Ai148;PVCre mice (n=4).

**Figure 4:**
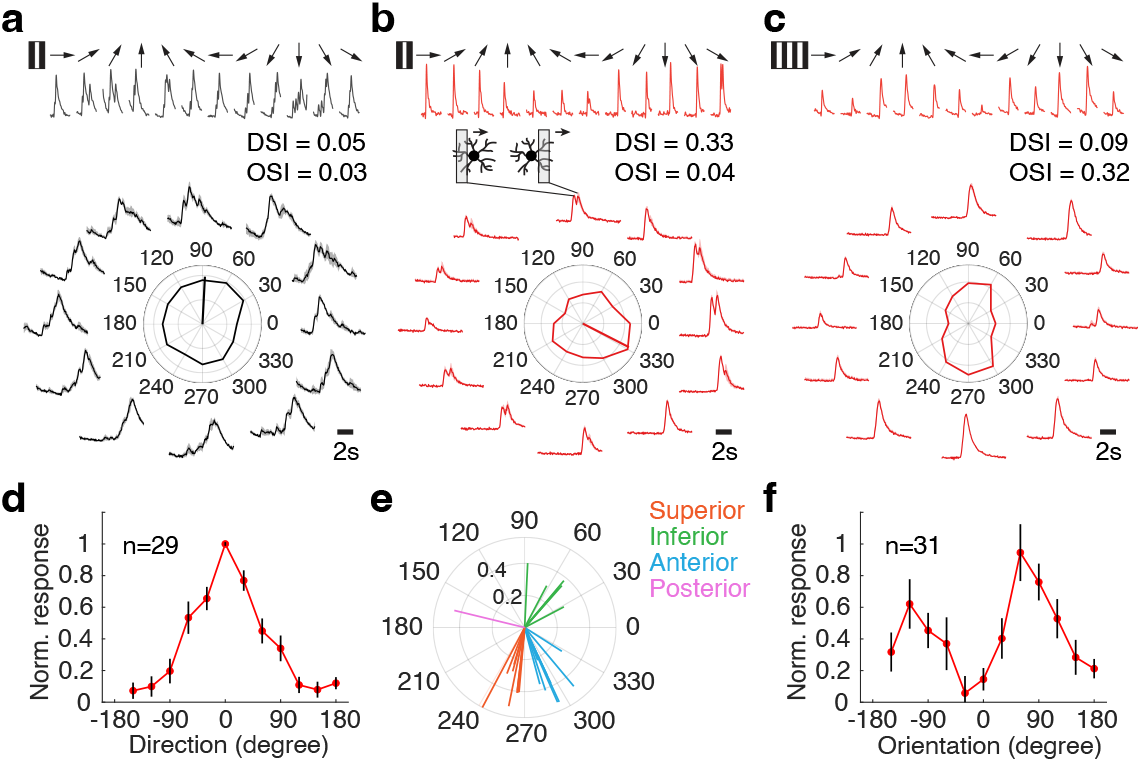
Direction and orientation selective responses of RGCs. **(a)** Response of a non-DS RGC. Top: A bright bar moving on a dark background along 12 directions. Bottom: Trial averaged calcium responses to different directions of bar movement. Gray shaded error bar is SD. The polar plot in the center shows normalized response and preferred direction of the RGC. DSI: Direction Selective Index; OSI: Orientation Selective Index. **(b)** Same as in (a), for an ON-OFF DS-RGC. Inset illustrates ON and OFF responses elicited by a bright bar entering and exiting the receptive field of the RGC. **(c)** Same as in (a), for an OS-RGC. **(d)** Mean tuning curve of DS-RGC population (n=29 RGCs). Black error bar is SD. **(e)** Preferred directions of ON-OFF DS-RGC subtypes (n=21 RGCs). **(f)** Mean tuning curve of OS-RGC population (n=31 RGCs). Black error bar is SD.

To identify orientation selective (OS-) RGCs, we measured calcium responses to drifting gratings presented at different orientations (Fig. 4c). Orientation selectivity was quantified by an orientation selective index (OSI) (see Methods). To distinguish OS-RGCs from DS-RGCs, we calculated the DSI of OS-RGCs from their responses to moving bars. A total of 31 RGCs which had OSI>0.3 and DSI<0.3, were classified as orientation selective (Fig. 4f). Note, it is possible some of these RGCs are bi-direction selective and not OS per se. However, OS cells have been described previously in the retina, while bi-direction selective cells have not.

### Calcium imaging at bipolar cell axon terminals

Finally, we tested whether the system’s resolution allows for imaging calcium activity at synaptic release sites. For this, we used retinas from Ai148;PCP2Cre transgenic mice^13^ that are known to express GCaMP6f primarily in type 6, type 2 and rod BCs^36,37^. Depolarization of BCs and subsequent glutamate release are associated with increased calcium influx at the axon terminal release sites^20^, making them ideal for imaging. Light-sheet excitation was targeted to the inner portion of the inner plexiform layer adjacent to the ganglion cell layer. Robust GCaMP6f expression was observed in the puncta, each ∼1-4 *μ*m, clustered across the imaging field, consistent with the size and density of release sites at BC terminals (Fig. 5a, b).

**Figure 5:**
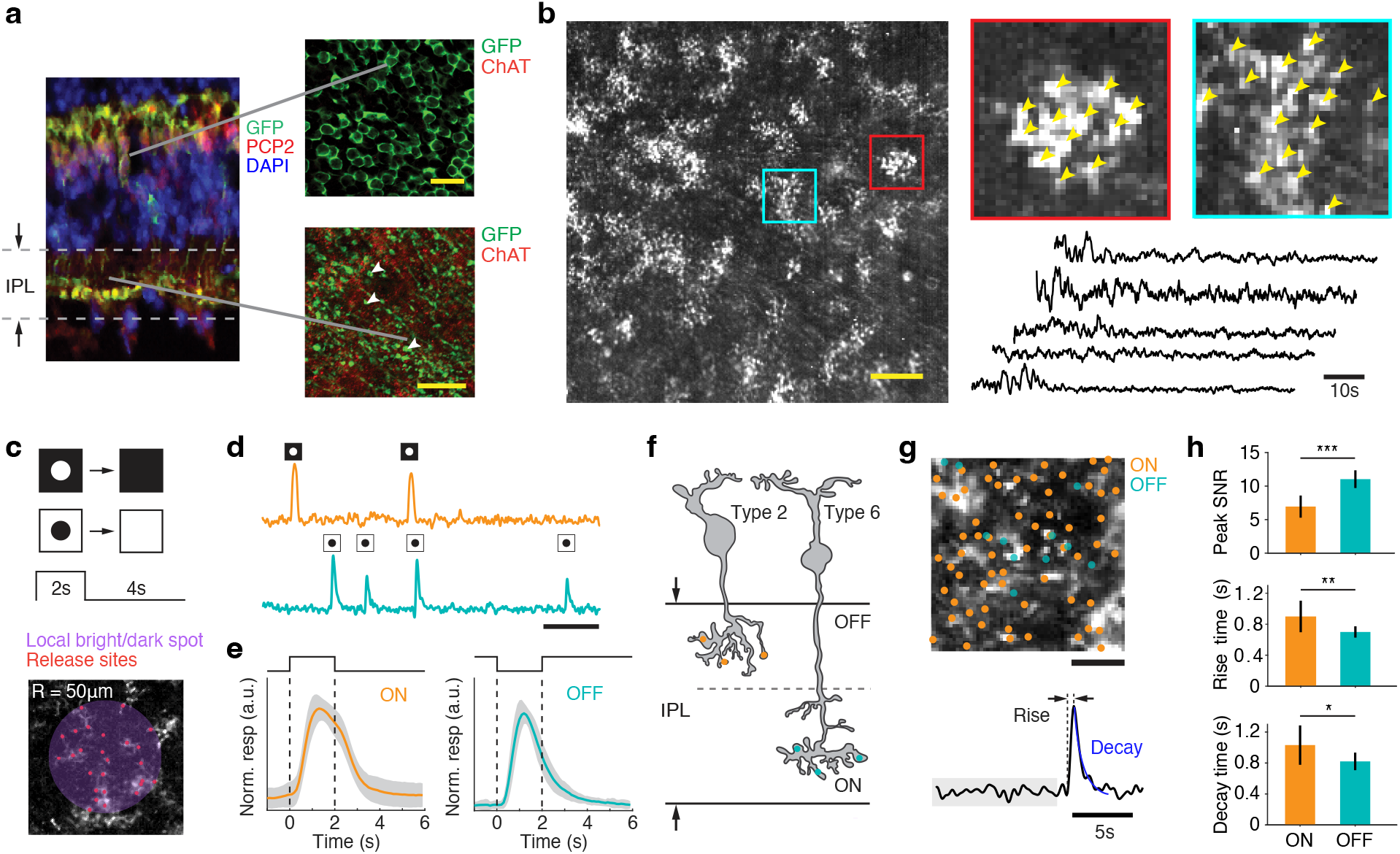
Calcium imaging of BC axon terminals. **(a)** Left: Colocalization of GFP (green) and PCP2 (red) showing soma and terminals of BCs in the retina of an Ai148;PCP2Cre mouse. DAPI: blue. Right: Flat-mount view of somata (top) and axon terminals (bottom). Scale bar: 20 *μ*m. **(b)** Left: SD projection image showing active release sites. Scale bar: 100 *μ*m. Right: Magnified view of regions indicated in red and cyan with spatial footprints (yellow arrows). Spontaneous responses (black traces) of 5 randomly selected release sites. **(c)** Top: 100 *μ*m diameter spots at 100% positive and negative contrasts, are presented for 2 s at n=34 locations. Bottom: Representative spot (shaded circle) and active sites (red dots). **(d)** Representative temporal fluorescent responses of a release site stimulated by a bright spot (top, orange), and a dark spot with 50 *μ*M L-AP4 (bottom, teal), appearing at these locations at random times (n=5 repeats for each location). Scale bar: 20 s. **(e)** Normalized calcium transients to bright (left) and dark (right) spots respectively, averaged over n=504 (ON, orange) and n=156 (OFF, teal) release sites. Dashed vertical lines indicate duration of the spot. Solid curve: mean. Shaded error bar: SD. **(f)** Schematized structure of a type 2 and a type 6 BC, and their axonal arbors in the inner plexiform layer (IPL). Orange and teal filled circles represent putative release sites in the OFF and the ON layers respectively. **(g)** Top: Release site locations on the SD projection image. Scale bar: 20 *μ*m. Bottom: Representative trace (black) of calcium transient used for estimating rise time, baseline fluctuations (gray) and time of decay (blue). **(h)** Ratio of peak value of calcium transient to the SD of baseline response (top), rise time (middle) and decay time constant from exponential fit (bottom), for ON (n=504, orange) and OFF (n=156, teal) release sites. Mean, SD and *P*-values with Bonferroni correction are: 6.92±1.72 (ON), 13.15±1.52 (OFF), *P* = 3.9 × 10^−7^ (top); 0.93±0.22 (ON), 0.71±0.10(OFF), 5.21 × 10^−3^ (middle); and 1.03±0.25 (ON), 0.82±0.11 (OFF), 3.1 × 10^−2^ (bottom). All OFF-responses in (d), (e), (g) and (h) were obtained with L-AP4.

In the absence of visual stimuli, the calcium fluorescence was relatively weak across the different release sites (Fig. 5b). To visually stimulate the BCs (via the photoreceptors), bright and dark spots were presented at different locations (Fig. 5c; see also Methods). A bright spot on a dark background induced strong calcium fluorescence transients across hundreds of release sites within a 730 *μ*m × 730 *μ*m field of view (Fig. b, d). Following disappearance of the spot, the fluorescence rapidly decayed to the baseline, indicating that these responses are likely produced by type-6 ON BCs (Fig. 5e, f). The release sites that exhibited a transient increase in calcium fluorescence in response to a dark spot on a bright background were sparser and had smaller response amplitudes compared to the ON release sites. Therefore, to unmask these responses, we blocked the ON pathway using L-AP4 – an mGluR6 agonist^38^. This significantly improved the signal to noise ratio (SNR) of the imaged responses, allowing unambiguous identification of these release sites (Fig. 5d, e, f). The responses rose and decayed sharply to the appearance and disappearance of the dark spot (Fig. 5d, e). In addition, moving the excitation and the imaging plane up toward the photoreceptors by ∼10 *μ*m improved the SNR of these release sites by ∼20%, indicating these responses were from type-2 OFF BCs (Fig. 5f). The temporal resolution and the SNR were sufficient for revealing the faster kinetics of the OFF responses in the outer layers of the IPL relative to the ON responses in the inner layers of the IPL (Fig. 5h), in agreement with previous studies^37,39^

## Discussion

We present a 1-photon light-sheet imaging system, that allows measurements of neural activity across large populations of retinal neurons at synaptic resolution, while simultaneously presenting visual stimuli to photoreceptors. The widefield planar sheet of light is confined to a layer ∼20 *μ*m thick, which allows for imaging calcium activity in large cohorts of neurons confined to specific layers in the retina. An axial separation is maintained between the excitation plane for fluorescence imaging and the focal plane for visual stimulation of the photoreceptors. Using this system, we were able to measure spontaneous and visual stimulus dependent responses of hundreds of RGCs routinely over a retinal area 1.5-2.25 mm^2^, corresponding to 50-80 degrees of visual angle in the mouse retina (Fig. 3a). The signal to noise ratio of the measured activity was sufficient to functionally classify RGCs in Ai148;PVCre retinas into 8 distinct types, consistent with previous morphological classification^11,12^. The resolution of the system also allowed measurements of calcium activity in individual synaptic release sites at BC axon terminals from Ai148;PCP2Cre retinas. We could distinguish the release sites associated with ON and OFF type BCs consistent with previous studies^13^.

Optical imaging techniques that allow monitoring ensemble activity of neurons within biological specimens, can offer new insights into how sensory signals are processed within specialized neural circuits. For example, 2-photon calcium fluorescence imaging^1,4,35^ has been widely used in both retinal and cortical research for several reasons: 1) It provides high spatial resolution; 2) infrared light scatters less than shorter wavelength light, which facilitates imaging deep inside the tissue; and 3) infrared light reduces the activation of photoreceptors when imaging retinal neurons. However, this approach requires a costly and complex pulsed femtosecond laser setup and provides data with either poor temporal resolution or limited scan areas due to point laser scanning.

Light-sheet fluorescence microscopy is an alternative imaging modality that utilizes planar illumination to optically section the sample, enabling rapid acquisition of images with high spatial resolution while minimizing photobleaching of the fluorescence indicator^40^. These advantages have led to an increased adoption of this technology for biomedical imaging, ranging from cultured tissues to *in vivo* imaging in small animals such as zebrafish and drosophila^41^. In particular, combining optical sectioning with synchronized delivery of excitation light^42^, allows capturing of subcellular dynamics in living cells as well as 3-D reconstruction of activity in living biological specimens^43^. However, constraints in delivering the planar excitation light orthogonal to the detection axis often require the sample to be embedded in agarose – a condition not ideal for *ex vivo* retina. This requires custom-designed systems that can accommodate the geometry of the sample for mounting and keep the sample viable for long-term imaging.

Light-sheet imaging allows robust measurements of calcium responses with high spatial resolution in *ex vivo* retina, and therefore offers several benefits. First, the ability to monitor neural activity at the resolution of single synapses across a large area will open the possibility of examining how excitatory and inhibitory synaptic inputs from genetically targeted interneurons are integrated over dendritic sub-compartments. This can offer new insights into circuit-specific computations^44^. Second, measurements of activity simultaneously in multiple genetically targeted cell types, combined with pharmacological or optogenetic manipulations^45^, can help distinguish the role of different presynaptic cell types in shaping the gain and nonlinearities of signal transfer. Third, measurements of large-scale neural activity in specific layers will enable characterization of how signal and noise correlations^46^ impact the encoding and transmission of information about visual features by different populations of neurons.

A challenge in retinal fluorescence imaging using 1-photon light-sheet is scattering of the excitation light. The 488 nm light scattered within the retina can potentially reach the photoreceptors and active the M-opsins in cone photoreceptors. Another challenge is contamination of fluorescence signals from neurites and other structures such as axon bundles expressing GCaMP (Fig. 3e). But, with the development of faster, brighter and long-wavelength sensitive indicators^7,47^, as well as soma and dendrite directed calcium indicators^48^, these challenges can be substantially mitigated. The architecture of 2-photon light-sheet imaging with the use of infrared lasers^43^ can be integrated into our framework to overcome limitations in spatial resolution. Our system also allows for multiplexing excitation light of different wavelengths that can be targeted to different retinal planes, and incorporating remote focusing and a tunable lens^49^ to perform rapid multi-plane imaging in the retina. Given the versatility, flexibility and high-throughput measurement capabilities of our imaging platform, we envision this system to become a powerful tool for large-scale interrogation of functional connectivity between cell types in the retina.

## Methods

### Animal and retina preparation

Procedures for animal care and use followed guidelines approved by the Institutional Animal Care and Use Committee at Duke University. The Ai148 floxed mice carrying the GCaMP6f transgene under the control of tetracycline transactivator tTA2 (TIT2L-GC6f-ICL-tTA2) (Jackson Laboratory, 030328), were crossed to (1) PVCre mice carrying a Cre allele in parvalbumin (PV) expressing neurons (Jackson Laboratory, 008069), and (2) PCP2Cre mice carrying Cre allele in Purkinje cell protein (PCP2) expressing neurons (Jackson Laboratory, 010536), to obtain the Ai148;PVCre and Ai148;PCP2Cre mice respectively. Mice (n=10 Ai148;PVCre and n=6 Ai148;PCP2Cre) with age between 2-10 months of both sexes were used for experiments. Retinas for experiments were obtained following previously established protocols^50^. Briefly, mice were kept under 12 hr light-dark cycle with *ad lib* access to food and water. For experiments, animals were dark-adapted for 12 hrs, then euthanized in complete darkness and under infrared illumination using infrared goggles. The eyes were enucleated, and retinas were dissected out in a petri dish containing sodium bicarbonate buffered Ames’ media (Sigma Aldrich, A1420) bubbled with 95% oxygen and 5% carbon dioxide (pH 7.4, temperature maintained at 33°C). A piece of retina ∼1.5 mm × 2 mm was cut from the ventral half and transferred to a custom-designed chamber containing oxygenated Ames’ solution. The chamber has a glass bottom for imaging and a glass side window for entry of light-sheet excitation. The retina was flattened gently using a hollow cylinder with a porous membrane (Spectra/Por RC dialysis membrane, 132677) that allows passage of solution and metabolites. The chamber containing the retina was transferred to the light-sheet microscope for imaging. Throughout the experiment, the retina was continuously superfused with oxygenated Ames’ solution (described above) with a gravity-fed perfusion system.

### Light sheet excitation

The excitation light was provided by a 488 nm laser (OBIS LX continuous wave laser, Coherent, Inc.). The laser beam was collimated using a fiber collimator (Thorlabs, Inc., F240APC-A) and expanded to 1 mm diameter by a pair of relay lenses with effective focal length 225 mm (Edmunds Optics, Inc., 47-365, 47-645). The expanded Gaussian beam is compressed by a cylindrical lens with focal length 50 mm (Edmund Optics, Inc., 33-228) and the resulting light-sheet is focused on the back aperture of the illumination objective (Mitutoyo, 5x, 0.14 NA, MY5X-802) by a pair of relay lenses of effective focal length 50 mm (Thorlabs, Inc., LBF254-050). A pair of orthogonal slits (Thorlabs, Inc., CP-20S, VA100C) controlled the lateral extent and the axial thickness of the light-sheet. The laser operating power was maintained between 0.1-15 mW, that produced 0.01-1.5 mW power at the sample. The depth of field was estimated by obtaining a stack of 200 images in 1 *μ*m steps, of 500 nm diameter fluorescent polystyrene beads (Spherotech, Inc., FICP-08-2) embedded in agarose placed inside a quartz cuvette (Thorlabs, Inc., CV10Q35F).

### Visual stimulation and calcium imaging

Visual stimuli were rendered using an OpenGL framework using custom written scripts in MATLAB (The Mathworks, Inc., Natick, MA). The stimulus patterns were streamed via an HDMI cable to the LightCrafter 4500 Digital Light Projector (DLP) (EKB Technologies, Ltd., DPM-E4500LUVBGMKII) and controlled by a custom GUI. The stimulus was displayed at 385 nm using a built-in LED, operated in the linear range. The display comprised of digital micromirrors arranged in a diamond pattern, with spatial resolution of 920 × 1040 pixels. To minimize spherical aberration of the projected stimulus image, a circular aperture (Thorlabs, Inc., SM1D12D) was placed in front of the DLP. The visual stimulus was collimated by a 100 mm aspheric converging lens (Thorlabs, Inc., AL50100) and the tube lens of Ti microscope (Nikon Instruments, Inc.). The final image was focused on the photoreceptors by a 10x Plan Fluor (Nikon Instruments Inc., MRH00101) or a 20x Super Plan Fluor (Nikon Instruments Inc., MRH08230) objective, rated to operate in the UV wavelength range, through the bottom glass surface of the chamber. The stimulus plane was offset from the imaging plane by controlling the distance between the DLP and the focusing lens (Fig. 1a). This offset allowed us to simultaneously use the same objective for visual stimulus delivery, and imaging calcium dependent fluorescence in RGCs and BCs. The following set of stimuli were used in our experiments: (1) bright bar (100% Michelson contrast) traveling along 12 different directions with a speed of 480 m/s, (2) grating (100% Michelson contrast) moving along 12 different directions with a speed ranging between 24-100 m/s, (3) full-field ‘chirp’ sequence comprising of dark (3 s), bright (3 s), contrast frequency modulation (0.5 - 8 Hz, 8 s period) and contrast amplitude modulation (0.5 - 2 Hz, 8 s period), repeated 7 times, (4) full-field light increment and decrements, (5) binary checkerboard patterns with checker size 10 - 50 *μ*m, and (6) local bright/dark spots (100 *μ*m diameter) repeated 5 times at each location manually selected from a template image. L-AP4 (L-(+)-2-Amino-4-phosphonobutyric acid, Tocris Bioscience, 0103) at 50 *μ*M was used to block mGluR6 receptors. The stimulus frames refreshed at 60 Hz. The stimulus brightness was calibrated using a photometer (Thorlabs, Inc., PM100D) and was set to ∼10^5^ P*/S-cone/s for experiments using neutral density filters (Thorlabs, Inc.).

Calcium images were captured by an ORCA Fusion camera (Hamamatsu Photonics) using the HC-Image software (Hamamatsu Photonics). GCaMP6f expression in RGC somas and BC terminals was reliably observed over a laminar depth of 20-30 *μ*m. A long-pass dichroic mirror (Thorlabs, Inc., DMLP425R) was used to reflect the UV stimulus and transmit the GCaMP6f emission. A long-wave-pass edge filter (Semrock, FF01-430/LP-25) and a GFP filter (Semrock, GFP-3035D) were placed after the dichroic to block UV light and allow emitted light, respectively, to reach the camera. Images are acquired at 10-50 Hz, at 16-bit resolution, with spatial binning of 2 or 4. Pixel size of projected image was calibrated for each imaging objective using a glass reticle with 10 *μ*m resolution. Calcium images were registered with visual stimuli by using timestamps from the camera and stored in the stimulus computer via a 6550-USB DAQ device (National Instruments, Corp.).

### Theoretical thickness of light-sheet

The thickness of the light-sheet determines the axial range over which the sample can be reliably imaged. The light-sheet produced by the excitation optics (Fig. 1a) has a Gaussian profile with a beam waist:

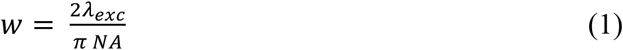

Here, *λ*_*exc*_ is the wavelength of excitation light and *NA* is the numerical aperture of the illumination objective. If *θ* is the half angle of the light cone generated by the objective and *n* is the refractive index of the medium between the sample and the objective, then Eq. 1 can be reformulated as:

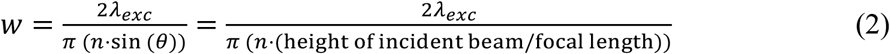

Given a 488 nm excitation wavelength, a ∼1 mm diameter beam, a 40 mm focal length illumination objective, and 1.33 refractive index of water, the beam waist is,

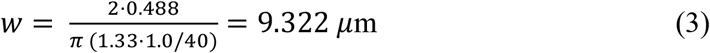

### Axial resolution and depth of field

The axial resolution of a microscope depends on the optical properties of the detection system and the refractive index of the sample. Considering the elongated shape of the intensity profile along the axial direction^51^, the theoretical axial resolution is the distance between the central maximum to the first minimum along the Z-axis:

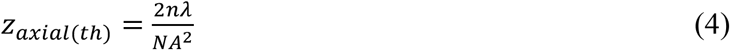

where *n* is the refractive index of the sample, *λ* is the wavelength of the emitted light and *NA* is the numerical aperture of the imaging objective. With 510 nm peak emission wavelength of GCaMP6f, *n* =1.38 for retina and 0.45 *NA* of imaging objective, the theoretical axial resolution is estimated as 6.95 *μ*m. The theoretical depth of field (DOF) is half the axial resolution:

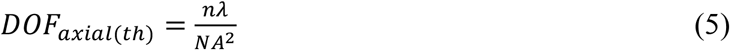

which is equal to 3.48 *μ*m.

Notably, these calculations assume that the detected light is emitted by a point source. However, scattering would tend to increase the Z-range of excitation, effectively increasing the axial resolution. Given the retina’s scattering coefficient, *μ*_*s*_, the effective axial resolution is given by the convolution of the expanded profile of the light-sheet, the excitation point-spread function and the detection point-spread function. To first order, the point spread functions can be approximated by Gaussians, which yields the following equation for effective axial resolution:

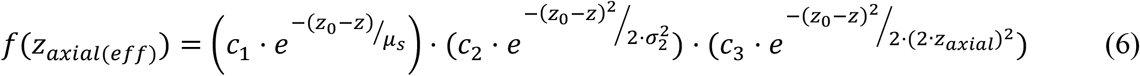

Here, *z*_0_ defines the plane of light-sheet and *σ*_2_ is the beam waist. This effective DOF, which is twice the *z*_*axial*(*eff*)_, was measured in the retina to be ∼22 *μ*m (Fig. 1f).

### Scattering of excitation light

The retina has a scattering coefficient of ∼57 cm^-1 17^, compared to ∼0.003 cm^-1^ of water. This means that the excitation light will undergo significant scattering as it travels through the retina. A large fraction of the incident photons is absorbed by the GCaMP6f protein, while the remaining fraction is scattered above and below the plane of excitation. The light scattered above the excitation plane can potentially reach the photoreceptors and activate them^22^, thereby producing visual responses independent of the stimulus.

To determine the degree of photoreceptor activation from scattered excitation light, we measured the total power of scattered light in the retina of an Ai148;PVCre mouse. Since scattered photons can travel along different directions, only a fraction of the scattered light reaches the photoreceptors. Therefore, to measure the intensity of scattered light that could activate photoreceptors, we used an aperture ∼2 mm × 2 mm roughly matching the size of the imaged retina (Supplementary Fig.1 a, b insets). The intensity of scattered light was converted to photoisomerization (P*) rate per cone expressing M- or S-opsin, using Baylor and Govardovskii nomograms^52,53^. Over a range of 0.1-3 mW laser power, the scattered light produced at most 10^4^ P*/M-cone/s (Supplementary Fig.1 a), and 1.0 P*/S-cone/s (Fig. 2c), which is ∼5 orders of magnitude lower than the photoisomerization rate produced by photopic UV stimulus^22^ used in our experiments (Supplementary Fig.1 b).

### Depth of field of imaging

Diffraction-limited point objects such as fluorescent beads are commonly used for assessing the spatial resolution of a microscope^16^. We used 500 nm diameter fluorescent beads coated with green-fluorescent dye embedded in 2% agarose gel (Fig. 1c top) for measuring the depth of field. Images were acquired every 1 *μ*m along the Z-axis, while keeping the Z-position of the sample and the excitation light-sheet unchanged. The (x, y) location of a bead was determined from the image with the bead in focus, and intensity was averaged over pixels within 1 standard deviation around the peak centered at (x, y). Using images at different Z-positions, an intensity profile of the bead was measured as a function of the axial distance (Fig. 1d). Since the detection objective collects more light from below the focal plane than above it, it leads to an asymmetrical intensity profile. Therefore, we symmetrized the intensity profile by reflecting the intensity curve about the excitation plane. By fitting a Gaussian function to the intensity profile and determining the full-width half maximum (FWHM) of the fit, we estimated the depth of field to be 8.5 ± 2.8 *μ*m, averaged over n=63 beads (Fig. 1d inset).

### Histology

Wholemount staining was performed on the retinas of Ai148;PVCre and Ai148;PCP2Cre mice. The retinas were fixed in 4% PFA for 45 minutes at room temperature and then incubated in 5% normal donkey serum (Jackson Immuno, C840D36) in 1X phosphate buffer saline (PBS) with azide (Santa Cruz Biotechnology, SC-296028) containing 0.5% Triton X-100 (Sigma Aldrich, 93443), overnight at 4C. The retinas were then incubated in primary antibodies on a rocker for 3-5 days at 4C, after which they were rinsed in 1X PBS and incubated in secondary antibodies overnight at 4C on a rocker. The retinas were then rinsed in 1X PBS, placed on a filter paper and mounted on a glass slide with sealed coverslips. For cryosections, after fixation the retinas were incubated in 30% sucrose/PBS for 4-5 hours, frozen in OCT (VWR, 25608-930) and then sectioned at 15-20 *μ*m thickness. The mounted retinas were imaged with a laser scanning confocal microscope (Nikon Instruments, Inc., Ti-2) using 20x/40x/60x air objective. The Z-stack of images were processed in FIJI software^54^, to identify laminae and GCaMP expressing cells.

The following primary antibodies were used: anti-GFP (1:1000, Rockland, 600-901-215), anti-ChAT (1:500, Millipore Sigma, AB144P), anti-PCP2 (1:500, Santa Cruz Biotechnology, Inc. sc-137064). Secondary antibodies conjugated to Alexa 488 (1:500; Invitrogen, A-11094), Alexa 555 (1:500; Invitrogen, A20187), and Alexa 647 (1:500; Invitrogen, A-21447), were each diluted at 1:500. DAPI (Molecular Probes, S36964) was used for nuclear staining.

### Active ROIs, calcium responses and inferred spikes

To eliminate noisy, out-of-focus structures near the boundary of the imaged retina, a rectangular area containing the active ROIs was selected. Non-uniform illumination was corrected by homomorphic filtering and stripe artifacts caused by scattering of excitation beam were removed by spatial high pass filtering^55^. Size of a template ROI was determined by first manually selecting contours around active somas (for RGCs) or active synapses (for BCs), and then estimating a mean radius from the ellipses fitted to the selected ROIs. The images were denoised by a Kalman Filter and were batch processed using the CNMF-E algorithm^25^ to extract the time-varying fluorescence traces,

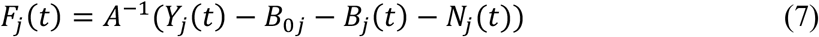

where *j* corresponds to the ROI index, *F*_*j*_(*t*) is the time varying fluorescent trace, *A* is the spatial matrix, *Y*_*j*_(*t*) is the raw trace, *B*_0*j*_ is the constant background, *B*_*j*_(*t*) is the time varying background and *N*_*j*_(*t*) is the time-varying noise. Manual verification was performed to remove overlapping ROIs and false positives. Steady drift in baseline fluorescence was corrected by subtracting the rolling 10^th^ quantile over a local time window from the raw trace. An estimate of Δ*F*(*t*)/*F*_0_ were obtained from the fluorescent trace prior to subtracting the baseline. The peak signal to noise ratio is given by the ratio of the average peak fluorescence of calcium transients to the standard deviation of the drift-subtracted baseline fluorescence:

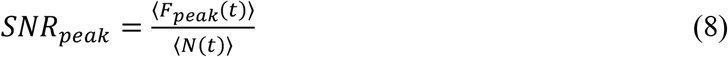

To infer spikes from temporal fluorescent traces, the calcium transients were first fit using an autoregressive model of order 1. The modeled transients were then deconvolved from the fluorescent trace to estimate spike count *s*_*j*_(*t*):

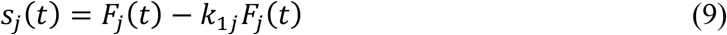

where, *j* corresponds to the *j*’th ROI, and *k*_1*j*_ corresponds to the first coefficient of the autoregressive model for the *j*’th ROI.

### Response clustering and RGC classification

To cluster RGCs into different groups, trial-averaged responses to the chirp stimulus were used. Principal components were calculated using the full ensemble of trial-averaged responses. The responses of each RGC were then projected onto the leading principal components that accounted for at least 80% of the variance (Fig. 3c). A Gaussian mixture model (GMM) with expectation maximization algorithm was fit to the projection values in the N-dimensional hyperspace with a pre-defined number of clusters determined from cross-validated Silhouette optimality test and Bayesian Information Criterion (BIC)^56^. Response clustering was tested with alternative methods: Hierarchical agglomerative clustering (HAC) and Spectral clustering (SC). (Supplementary Fig. 2). Since each functionally distinct RGC type tiles the retinal space^28^, the nearest-neighbor distance (NND) was computed for each cluster of RGCs (Fig. 3 g-h). Spike counts were obtained by the method described above (Eq. 9), and the number of spikes corresponding to the count was uniformly distributed across the bin to obtain spike times. Trial-to-trial variability was estimated using both calcium responses and inferred spike times, for different groups of RGCs (Supplementary Fig. 3d).

### Optimality and accuracy of clustering

To determine the quality of GMM fit to the projection values, we estimated Bayesian Information Criterion (BIC) values for different cluster sizes. The BIC value is defined as:

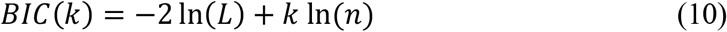

*L* is the maximum of Bayes likelihood, *k* is the number of estimated clusters and *n* is the number neurons. Since HAC and SC do not rely on model fits, optimality of cluster size was assessed by estimating the Silhouette coefficient for different cluster sizes using different clustering methods.

For each cluster size, the Silhouette coefficient *S*(*i*) for the *i*’th point in a cluster was calculated as:

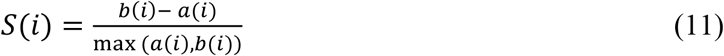

where, *a*(*i*) is the mean distance from the *i*’th point to all other points in the cluster and *b*(*i*) is the minimum of all distances from the *i*’th point to all other points in all other clusters. Normalized peri-stimulus time histograms of fluorescent traces were used for clustering and Silhouette coefficients were calculated for n=100 iterations for each cluster size. The cluster size with the highest ratio of median to absolute median deviation was chosen as optimal (Supplementary Fig.2). For GMM, both the Silhouette coefficient and Bayesian Information Criterion for model fits were used to cross-validate the optimal cluster size.

To determine clustering accuracy, responses were first clustered using three different clustering methods: GMM, HAC and SC, using a pre-defined cluster size. The median population temporal response for each cluster was then used to calculate pairwise correlation between responses, for each clustering method. Groups from each clustering method, with the highest pairwise response correlation, were assigned to the same functional type. The nearest-neighbor distances between RGCs for each group were assayed to confirm that the RGCs belonged to a unique functional type.

### Quantification of stimulus preference

Directional preference was quantified by the Direction Selective Index (DSI):

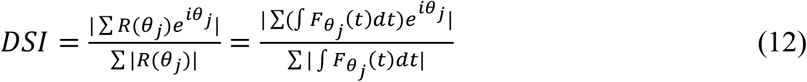

where *R*(*θ*_*j*_) corresponds to the area under calcium fluorescence response curve 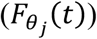 for a 100% contrast bright bar on a dark background moving along the direction *θ*_*j*_. RGCs with DSI>0.3 were selected as DS-RGCs^34^. ON-OFF DS-RGCs were identified by two response peaks, corresponding to the ON and OFF responses to the entry and exit of the bar over the receptive field (Fig. 4b).

Orientation preference was quantified by the Orientation Selective Index (OSI):

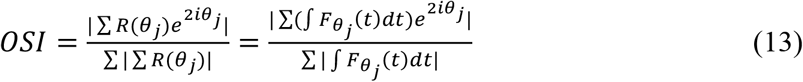

where *R*(*θ*_*j*_) corresponds to the area under calcium fluorescence response curve 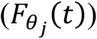 for a 100% contrast grating oriented along *θ*_*j*_. All RGCs with OSI>0.3 and DSI<0.3 were selected as OS-RGCs^34^. The grating moved in 12 directions at a speed of 24 *μ*m/sec, repeated 5 times.

### Receptive field estimation

Black and white checkerboard patterns with checker size ranging between 15-50 *μ*m, refreshing at 60 Hz, were used to characterize spatiotemporal receptive field (RF) of RGCs. Temporal calcium traces were low pass filtered, and then spikes were inferred using methods described above (Eq. 9). The sequence of checkerboard images *I*(*x, y*) preceding each spike *s*(*t*_*j*_) was weighted by the inferred spike count and averaged over the number of spike events to obtain the spatiotemporal RF.

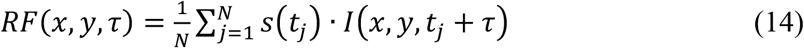

Here, *N* is the total number of spikes and *τ* is the time lag between a spike and a preceding image.

Calcium transients have a decay time of 300-400 ms^7^, therefore images presented over a 300 ms window preceding a spike event (i.e., *τ*<0.3 s), were used for estimating the temporal RF. The mean spatial image at each time lag was spatially filtered with a Gaussian of standard deviation 25 *μ*m. The pixel values from the spatially filtered image within a 300 *μ*m window centered around the RGC soma: (*c*_*x*_, *c*_*y*_) were averaged at each time lag to obtain the temporal RF.

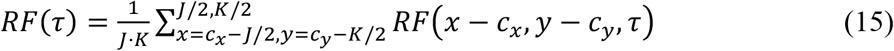

The temporal RF was fit with a parametric function *g*(*t*)

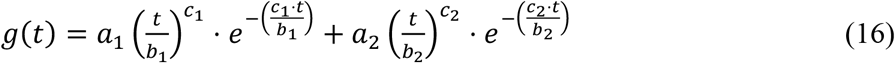

and the mean image corresponding to the closest peak (for ON RGC) or trough (for OFF RGC) to the spike event was used as the representative 2-D spatial RF. The RF center was fitted with a two-dimensional Gaussian with 1 SD boundary around the center maximum or minimum.

### BC response characterization

For extracting the ROIs for each release site, the following threshold values were used: (1) peak signal to noise ratio (SNR_peak_) = 2, (2) spatiotemporal pixel intensity correlation = 0.7, and (3) ROI template diameter = 1-4 *μ*m. After ROI extraction, duplicate and overlapping ROIs were manually removed. Fluorescence transients associated with the spot stimulus were identified by using a threshold value 5 times the median absolute deviation (MAD) of the entire temporal response. The peak signal to noise ratio, rise time, and decay time were used to characterize the kinetics of the response transients. The time constant for decay was obtained from an exponential fit to the temporal response curve within a 4 s window following the response peak. Positive (negative) contrast preference was determined by an increase followed by a decrease in calcium fluorescence to the appearance followed by disappearance of a bright (dark) spot.

### Statistical analysis

Statistical analyses of data were done using custom scripts written in MATLAB (Mathworks, Natick, MA) and CAIMAN codebase^25^. Summary data are presented as mean ± SEM, mean ± SD, or median ± MAD, as noted in figure legends or text. Statistical significance was determined from P-values with appropriate corrections for multiple samples of different sizes.

Optimality and reliability of clustering were determined using K-means distance, Silhouette Coefficient and Bayesian Information Criterion.

## Acknowledgements

This work was supported by NIH Brain Initiative Grant 1R34NS111645-01 (G.D.F.) and NIH K99EY032119 award (S.R.). We thank members of the Field lab for comments on the manuscript.

## Author contributions

S.R., A.S., and G.D.F. conceived the study. S.R., D.W. and Y.G. designed the microscope.

A.M.R. helped design and test the retina chamber. S.R. calibrated the imaging system, performed the experiments, and developed a pipeline for image analysis. S.R. and B.P. analyzed the data.

M.T. and M.L.S. assisted with mouse genetics and planning. S.R. and G.D.F. wrote the manuscript.

A.S. and Y.G. helped edit the manuscript.

---

**Supplementary Figure 1:**
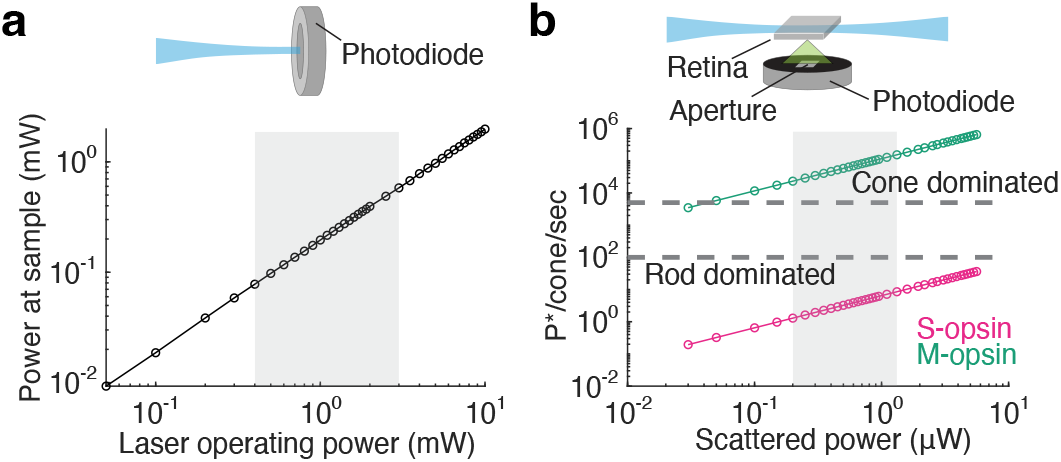
Photobleaching from scattered excitation light. **(a)** Top: Schematized setup for measuring power at the sample location using a photodiode. Bottom: Measured power versus operating power of the laser. Gray shaded region shows the range of laser power used in a typical experiment. **(b)** Top: Schematic showing power of scattered light from the retina of Ai148;PVCre mouse, measured by a photodiode through a 2×2 mm^2^ aperture. Bottom: Photoisomerization rate for M- and S-opsin cones as a function of the measure power of scattered light. Gray shaded region same as in (a).

**Supplementary Figure 2:**
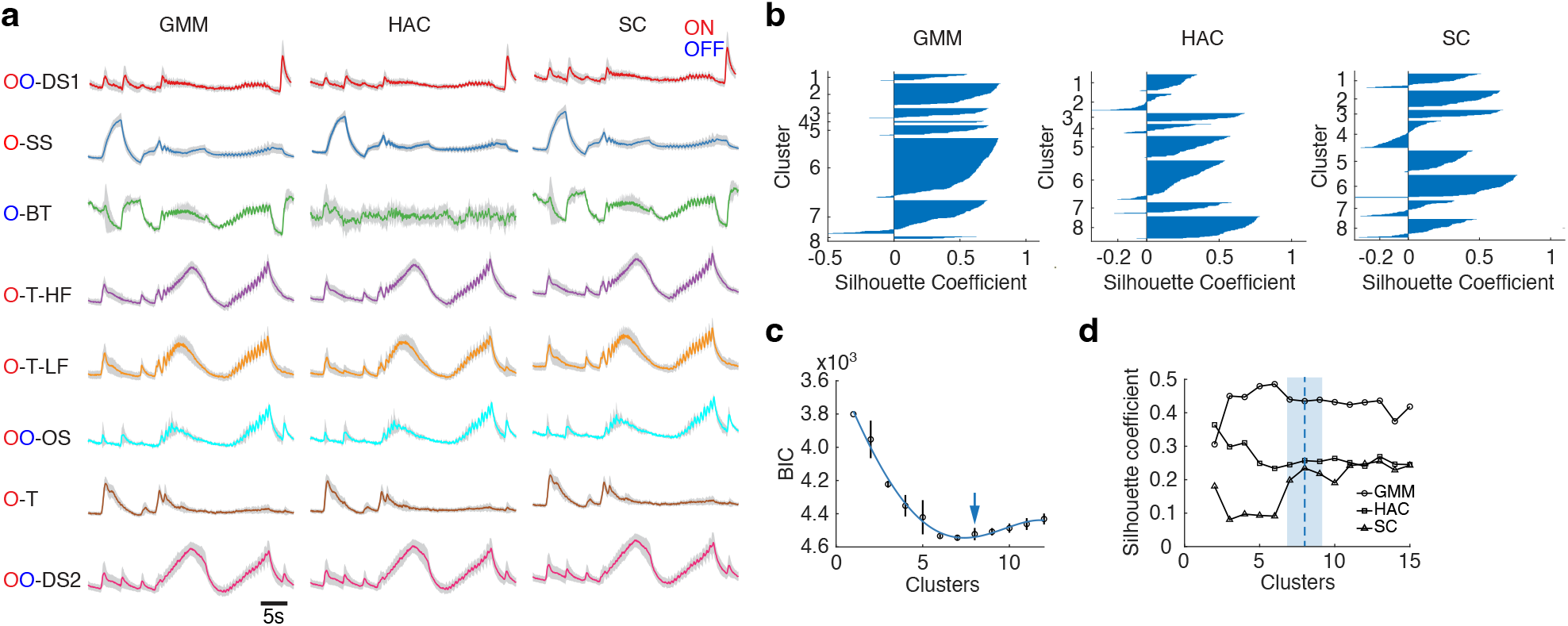
Comparison of response clustering using different algorithms. **(a)** RGCs clustered into distinct functional types based on clustering of responses to the chirp stimulus using: (1) Gaussian mixture model (GMM), (2) Hierarchical agglomerative clustering (HAC) and (3) Spectral clustering (SC). Results are representative of data shown in Fig. 3. ON: O (red), OFF: O (blue); DS: direction selective; SS: slow sustained; BT: brisk transient; T-HF: transient high frequency; T-LF: transient low frequency; OS: orientation selective; T: transient. Solid colored line: Mean population response. Gray shaded error bar: SD. **(b)** Silhouette test of optimality of cluster size for GMM, HAC and SC clustering methods. Shaded area shows silhouette coefficient (see Methods) for each cluster size. **(c)** Bayesian Information Criterion (BIC) for different number of clusters applied to the GMM fit. Black circles: Criterion values. Black vertical line: SD from n=100 iterations of model fit. Blue solid line: Polynomial fit. Blue arrow: Optimal number of clusters that distinguishes different response types. **(d)** Mean Silhouette coefficient for different number of clusters from each clustering method, GMM, HAC and SC, each indicated by a different marker type. Blue shaded region: Approximate range of cluster size with highest coefficient values across the three clustering methods. Dashed vertical line: Optimal cluster size.

**Supplementary Figure 3:**
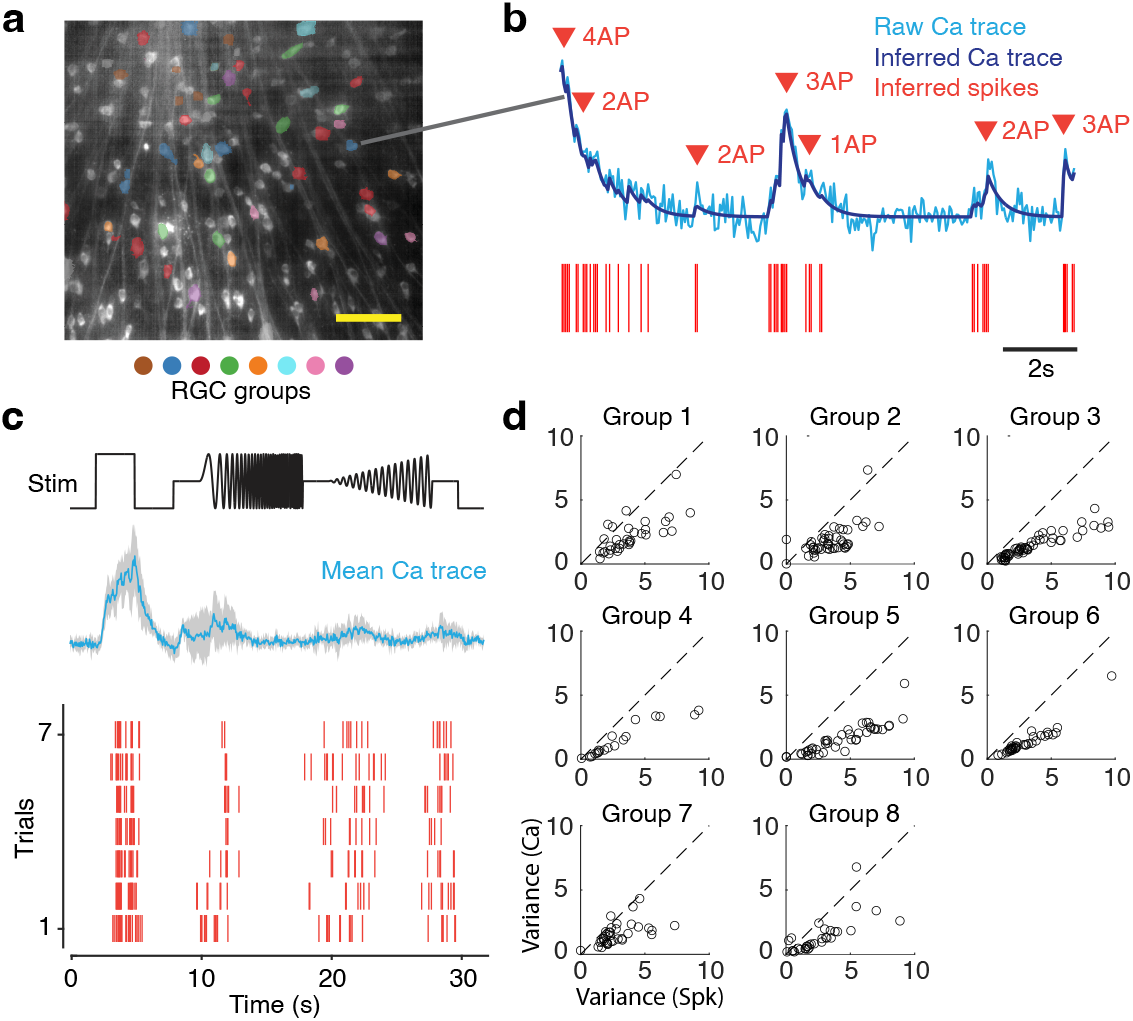
Statistics of inferred spikes of RGCs. **(a)** Median projection image showing RGCs belonging to different types (colored circles). Scale bar: 100 *μ*m. **(b)** Spikes inferred from deconvolved filtered fluorescence trace (dark blue). Unfiltered trace in light blue and inferred spikes in red. The number and timing of action potentials (APs) are indicated by red arrows. **(c)** Top: Chirp stimulus trace (black). Middle: Trial averaged (n_trial_=7) calcium responses of a representative RGC (solid blue line). Shaded error bar: SD. Bottom: Inferred spike times across trials. **(d)** Trial-to-trial variance of calcium responses and inferred spikes for each type of RGCs (n=8 types). Black circles: Individual RGCs.

